# Mutation of a palmitoyltransferase ZDHHC18-like gene is responsible for the minute wing mutation in the silkworm (*Bombyx mori*)

**DOI:** 10.1101/101618

**Authors:** Ye Yu, Xiaojing Liu, Xiao Ma, Zhongjie Zhang, Na Liu, Chengxiang Hou, Muwang Li

## Abstract

Pulpal- and adult-stage minute wing (*mw*) mutants of the silkworm *Bombyx mori* have much smaller wings than those of the wild type. Herein, we report the genetic and molecular mechanisms underling the formation of the minute wing. Fine mapping and positional cloning revealed that a palmitoyltransferase *ZDHHC18-like* gene (*BmAPP*）was responsible for the *mw* mutation. CRISPR/Cas9 screening of the *BmAPP* gene in wild-type embryos revealed that a significant proportion of adults had the *mw* mutation. In an *mw* mutant strain, u11, a 10-bp insertion in the promoter of a novel gene resulted in low-level translation of a protein belonging to the DHHC family. A PiggyBac-based transgenic analysis showed that the promoter induced the expression of a*BmAPP* gene promoter in the wild type but not the *mw* mutant. These findings indicate that functional deletion of this gene promoter accounts for the *mw* mutation in silkworm. The possibility that *BmAPP* gene is involved in Hippo signalling pathway, an evolutionarily conserved signaling pathway that controls organ size, is discussed.

## 1. Introduction

Insects are the only winged invertebrates among arthropods. They are also the most numerous and most widely distributed animal group(Trivers and Hare 1976). The ability to fly is a key factor underlying their expansion, invasion, proliferation, and reproduction(Engel and Grimaldi 2004). In *Drosophila*, the imaginal discs develop in the embryo and eventually form all the adult fly structures, such as the wings, halteres, and legs(Neto-Silva, Wells et al. 2009). The *Drosophila* wing has become an important model system in evolutionary biology, ecology, physiology, and other fields(Zera and Denno 1997), McCulloch, Wallis et al. 2009). The wings of insects can evolve in response to a changing environment(Lewin 1985). Recent research has revealed the roles of genetic, hormonal, and environmental control in molecular mechanisms underlying insect wing mutations(Roff 1994). Therefore, wing mutations appear to be an adaptation mechanism in the process of insect evolution.

*Drosophila* has been used as a model insect in genetic research. Sharma et al. previously reported that the *wingless* gene was responsible for wingless *Drosophila*(Sharma and Chopra 1976), Cabrera, Alonso et al. 1987). Morata and Garcia-Bellido (1976) discovered that a mutation in the *Ubx* gene led to poiser change in the wing(Garcia-Bellido, Ripoll et al. 1976). Previous studies reported that the vestige wing (*vg*) gene (Williams, Scott et al. 1990) gave rise to a trace wing mutation and that another gene, *blisters*, caused bullate wing(Brower and Jaffe 1989), Zusman, Patel-King et al. 1990, Brabant and Brower 1993, Prout, Damania et al. 1997. The findings of these studies are useful for investigations of other genes that may be involved in insect wing formation.

The silkworm *Bombyx mori* is an important lepidopteran model organism. A long period of artificial selection has produced *B. mori* with wing degeneration, as well as *B. mori* with and without the ability to fly, unlike wild-type *B. mori*, all of which can fly. Gene mutations associated with wing development in *B. mori* result in different shaped wings or abnormal colored wings. Thus far, 20 different *B. mori* wing mutations have been described. These include aplerism, Crayfish pupa, *vg*, minute wing (*mw*), and winglet (*rw*). Many studies have investigated *B. mori* wing mutations. Among those, the molecular mechanisms underlying the *fl* (Fujiwara and Hojyo 1997), Matsunaga and Fujiwara 2002 and non-lepis wing (*nlw*) (Zhou, Tang et al. 2004), (Zhou, Li et al. 2006) mutations are well known, but the mechanism underlying other *B. mori* wing mutations has not been elucidated.

The silkworm *mw* gene mutation is controlled by a recessive gene. The *mw* silkworm gene has been mapped to the 22nd linkage group. In *mw* mutants, the wings of the pupal and adult stages are much smaller than those of the wild type (Fig. 1). In a previous study, we constructed a linkage map of *mw*. As shown by a BLAST search of the *B. mori* genome database, the STS T2272 marker is located at the site of 1.265 Mb in nscaf3056. The *mw* gene is located between the end of the nscaf3056 and T2272, which has 275 kb(Ma, et al., 2013). In the present study, we analyzed all 15 candidate genes located in the 275 kb region. In addition, we characterized a 1500-bp promoter of *B. mori mw* and conducted a PiggyBac-based transgenic analysis.

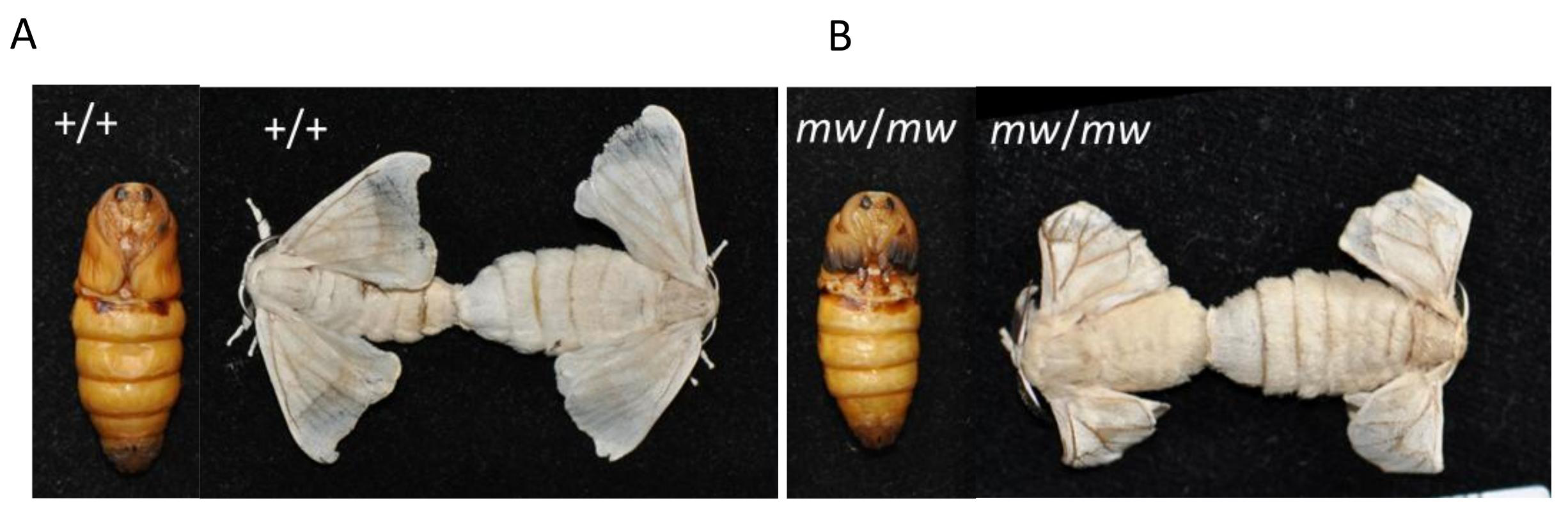
Phenotypes in the pupa and moth of the wild type and mw mutant strains. (A): The pupa and moth of the p50 strain (+/+). (B): The pupa and moth of the mw/mw mutant strains u11.*+/+* homozygote p50 strains, with bigger wings in the different periods. *mw/mw* homozygote u11 strains, with smaller, slightly curled wings that did not cover the metathorax and hind wings in the pupal and adult stages.

The results revealed that the palmitoyltransferase ZDHHC18-like (*BmAPP*) gene was responsible for the *mw* mutant and that the mutation was caused by a 10-bp insertion in the promoter region. They also showed that the promoter induced the expression of the reporter in the wild type but not in the *mw* mutant. These results indicate that functional insertion of the *mw* promotor is responsible for the *mw* mutation in silkworm. These findings may be useful in future lepidopteran pest control.

## Results

### Mapping the *mw* locus of *B. mori*

A linkage map for *mw*, which was located on chromosome 22, was constructed based on BC_1_M progenies. The BC_1_M progenies were obtained by crossing u11 females (*mw/mw*) with (u11×p50) F1 males (*+/mw*). Linkage maps were generated by MAPMAKER 3.0 using the Kosambi function, as described elsewhere (Lander, Green et al. 1987). First, we developed five STS markers (T2238, T2210, T2269, T2298, and T2272) on nscaf3056 of chromosome 22 (Fig. 2A). Finally, the *mw* locus was mapped to the 275 kb region between marker T2272 and the end of nscaf3056 (Fig. 2B). Within this region, 15 candidate genes were predicted according to the SilkDB database (Xia, Cheng et al. 2007) (Fig. 2C).

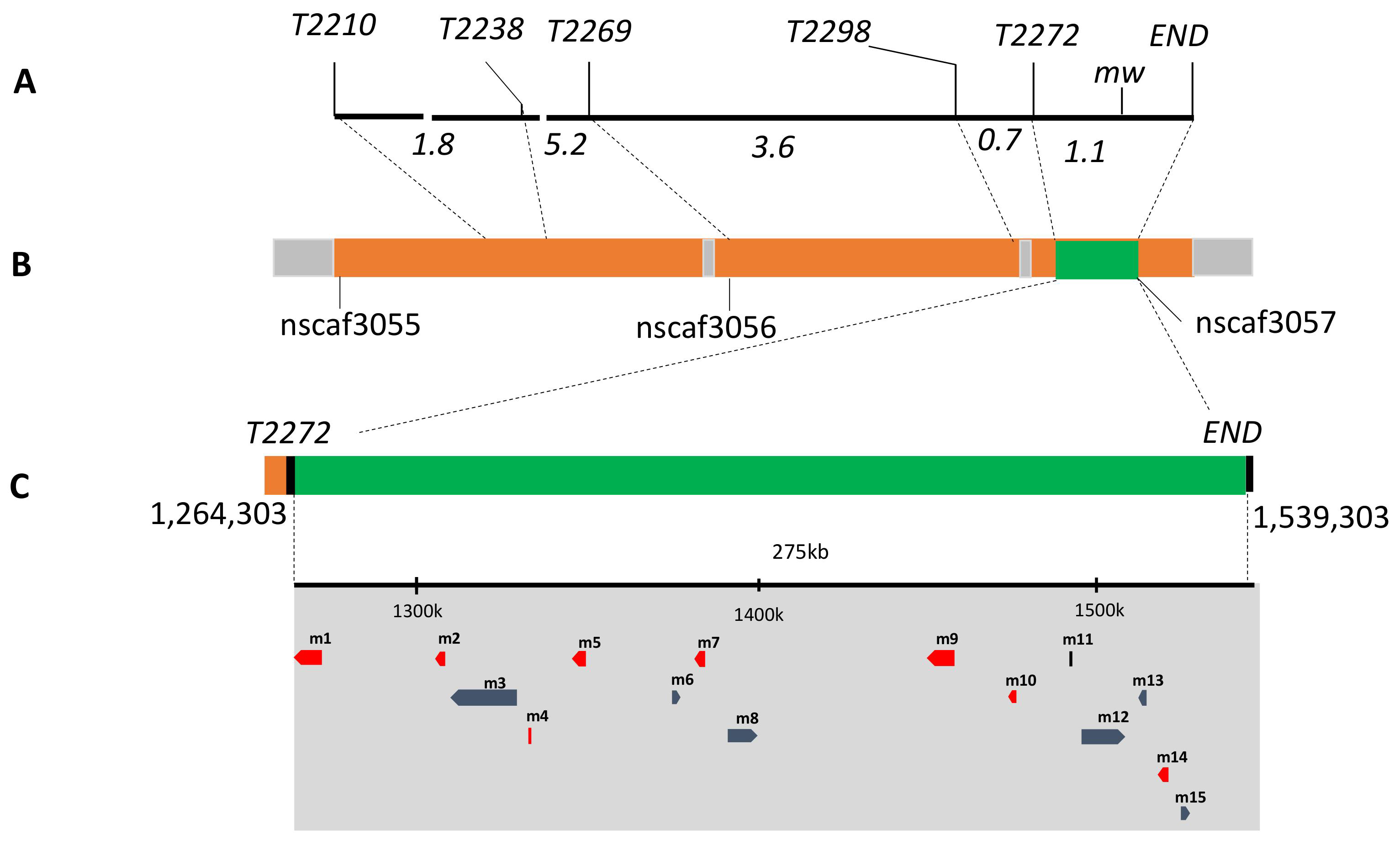
mw mapping on Bombyx mori chromosome 22 and identification of candidate genes. (A)A linkage map based on BC1M genotyping was constructed for group 22. Five sequence-tagged site (STS) markers and the mw locus are shown above the map. Distances between loci (cM) are shown below the map. (B)Scaffold map of chromosome 22, based on data from the SilkDB database. Orange boxes represent the assembled scaffolds, the names of which are given beneath. The STS markers used for genotyping are localized on the scaffold positions, as indicated by the dotted lines. The green box shows the candidate region linked to the mw locus. The enlarged candidate region is shown beneath along with the position on nscaf3056 of T2272 and terminal, which fix the candidate region. (C)Above is shown the coordinates of nscaf3056. The gray area represents the candidate region, which includes 15 predicted genes according to SilkDB. For convenience, they are named m1 to m15 based on their position. The red boxes show the eight genes that were excluded as candidates for the reasons described in the results section. The blue boxes indicate the candidate *mw* mutant genes. The directions of the transcription of these genes are represented by arrowheads.

### Identification of the *mw* candidate gene

Previous studies demonstrated that the phenotype of *mw* mutants was minute wings and that *mw* was expressed in the wing disc. Therefore, we examined the mRNA expression levels of 15 candidate genes in the wing discs of the wild-type P50 strain and *mw* mutant u11 strain (Fig. 3). Eight genes (indicated in red in Fig. 2B) were excluded because their mRNAs were not detected in the wing discs in 30 RT-PCR cycles. The other seven candidate genes (indicated in blue in (Fig. 2B) were expressed in the wing discs. Among these, the mRNA expression level of gene No.15 was much lower in the *mw* mutant u11 strain compared with that in the wild-type P50 strain. The expression levels of the other six candidate genes were identical in the wild-type and *mw* mutants (Fig. 3), and the cDNA sequences of these six candidate genes were the same in the u11 and p50 strains. A subsequent analysis of the mRNA expression level of candidate gene No. 15 in the wild-type p50 strain and *mw* mutant strain u11 of wandering-stage larvae suggested that candidate gene No. 15 was likely responsible for the *mw* mutation.

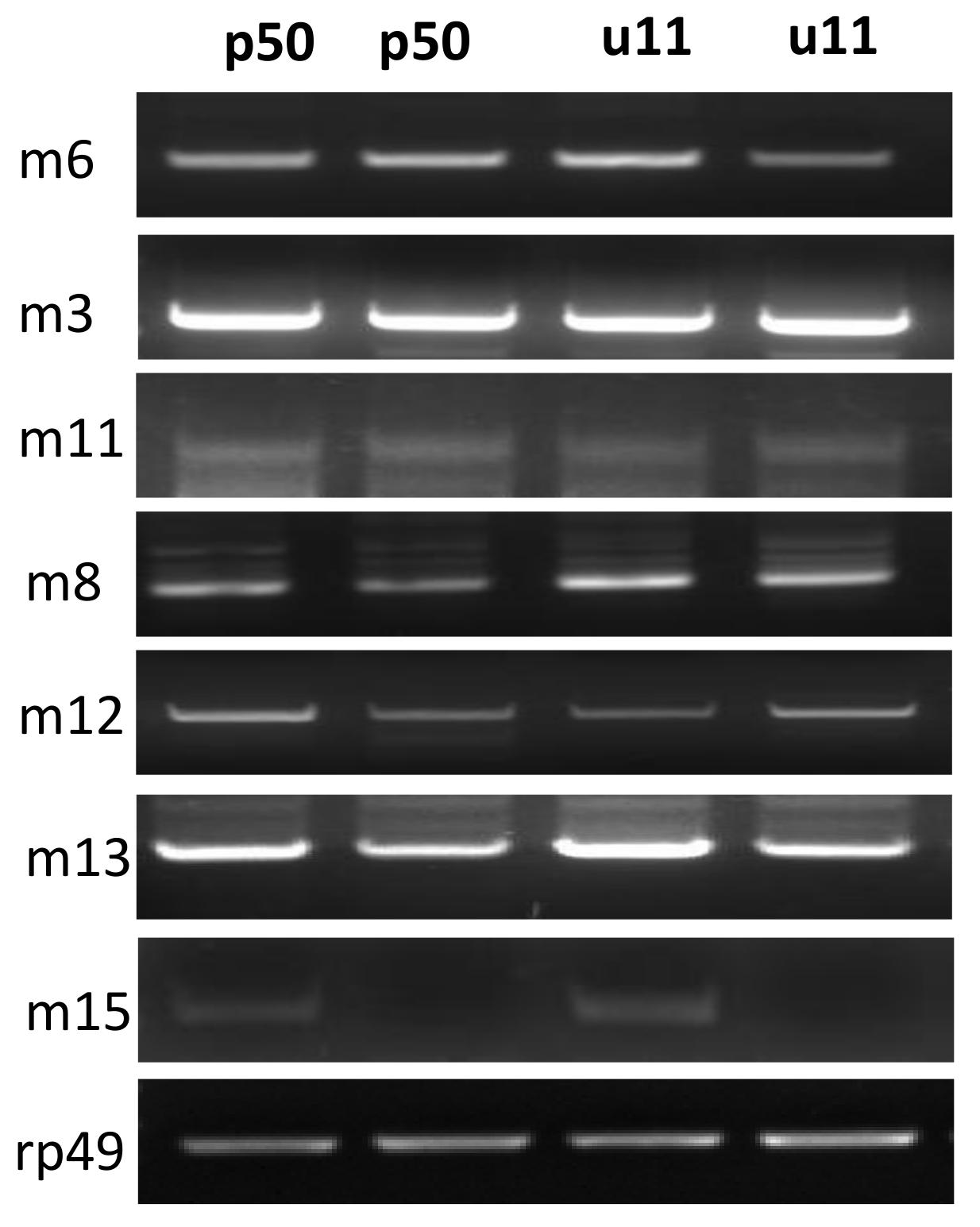
Semiquantitative RT-PCR analysis of the mw candidate genes. +/+ denotes p50, and mw/mw refers to u11. There were two repetitions. Candidate genes numbers m3, m6, m8, m11,m12, m13, and m15 are shown above. Total RNAs from the wing discs of wandering-stage wild-type p50 and mw mutant strains were used for the an0alysis. The primers used for RT-PCR are listed in Table 1. RP49was used as an internal control.

### CRISPR/Cas9 verification

To verify whether candidate gene No.15 was responsible for the minute wing mutation in strain u11, CRISPR/Cas9 was performed to suppress the expression of the corresponding gene in Nistari. As an emerging genome editing technology, CRISPR/Cas9 has been used for genetic analyses in many animal studies, including studies of the lepidopteron model insect *B.mori*(Dong, Chen et al. 2016). Figure 4A shows the genomic structure of *Bm-mw*, with two exons and an intron. Screening of the ORF of *Bm-mw* according to the GGN19GG rule (Hruscha, Krawitz et al. 2013) identified S1 and S2 sgRNA target sequences (Fig. 4A). S1 and S2 were located on exon 1 and exon 2, respectively, and the fragment spanning the two sites was 1486 bp in length (Fig. 4).

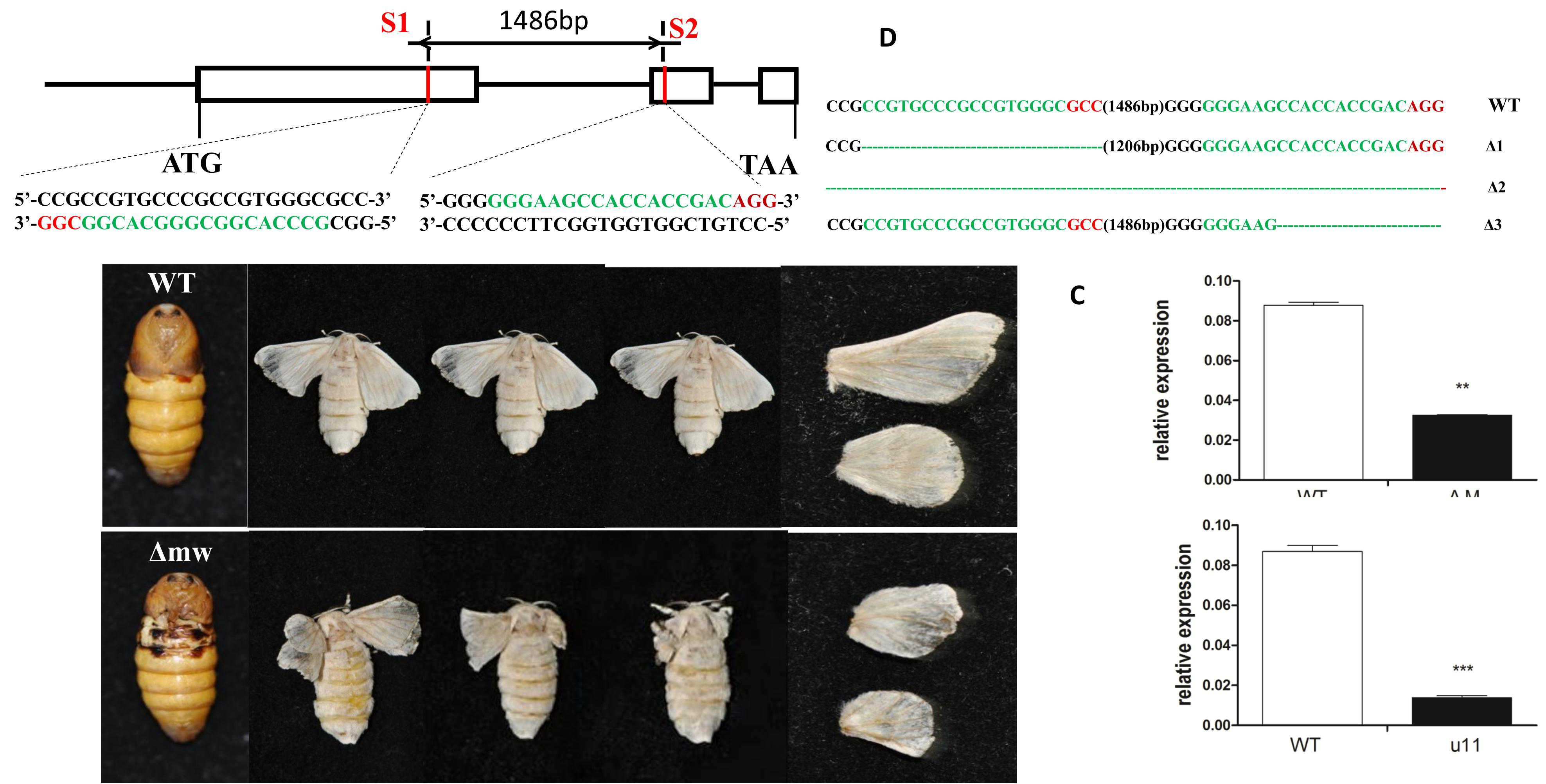
Schematic diagram of sgRNA targeting sites. The two boxes indicate the two exons of *mw*, and the black line represents the gene locus. The sgRNA targeting sites S1 and S2 are located on the sense strand of exon-1 and the sense strand of exon-2, respectively. The sgRNA targeting sequence is shown in black, and the protospacer adjacent motif (PAM) sequence is shown in red. (B)–Mutants with abnormal wings induced by Cas9–sgRNA injection. Phenotypes of the adults injected with sgRNA/Cas9 of the *mw* candidate gene m15 (right) at the embryonic stage and wild-type(left). (C) Various deletion genotypes in G0-injected embryos. The numbers in brackets in the middle of each sequence refer to the 1486-bp-long interspace fragment that was found between the S1 and S2 sites. The red sequence indicates the PAM sequence.

The sgRNAs that were synthesized in vitro were mixed with Cas9 mRNA and injected into preblastoderm embryos. In total, 480 eggs were injected with the Cas9/*mw*-specific sgRNA mixture at a concentration of 300 ng/μL Cas9 and 300 ng/μL sgRNA. Seventy-eight eggs hatched normally, and the hatching ratio was 16.25%. The hatching ratio in the controls (*N* = 480) was 58.75%. In the embryos that were injected with specific sgRNA and Cas9, approximately 50% showed wing defects. The control group showed normal phenotypes(Figure 4B). The majority of the mutants exhibited minute wings and curly wings. The adults injected with sgRNA and Cas9 exhibited malformed, curly wings, and the forewings did not cover the hind wings. The pupa of the u11 strain exhibited a similar phenotype, with the minute wings not covering the metathorax in the pupal stage (Fig. 4B).

To confirm that the phenotypic defects described above were due to genomic mutagenesis induced by injection of Cas9/sgRNA targeting *Bm-mw*, genomic DNA was extracted from five randomly selected silkworm mutants. The fragments spanning one or both target sites were then amplified, sequenced, and subjected to a PCR-based analysis. Both insertion and deletion mutations were detected in all five individuals (Fig. 4C). The maximal length of the deletion between S1 and S2 was approximately 300 bp. The transcript level of -mw is down-regulated significantly in mutants with sgRNA and u11(Figure 4D). The mutations were detected in different targeting sites. CRISPR/Cas9 injection showed that mutants with candidate gene No.15 developed minute wings similar to those of strain u11, suggesting that candidate gene No.15 was the key gene responsible for minute wings in the u11 strain.

### Defects of*Bm-mw* in the u11 mutants

According to the annotation, candidate gene 15 represented *Bm-mw*, a previously cloned silkworm gene (GeneID:101745665) that encodes a palmitoyltransferase ZDHHC18-like gene. To clarify the structure of *Bm-mw* in the u11 mutants, sequences of *Bm-mw* in the wild-type p50 were obtained from the NCBI database. *Bm*-*mw* consisted of three exons encoding a protein of 110 amino acid residues and spanned approximately 2.8 kb on nscaf3056 (Fig. 5A). To detect the defects caused by *mw* in the u11 mutants, the genomic sequences of *Bm-mw* in the u11 strain were compared with those of the wild-type p50 strain. PCR amplification was carried out using u11 and p50 genomic DNA as the template. In the *mw* mutant u11 strain, three exon sequences were undifferentiated as compared with those of the wild-type p50 strain. A RT-PCR analysis revealed no expression signal of *Bm-mw* in the u11 mutants and a 10-bp insertion before the initiation codon (Fig. 5B). This insertion may adversely affect the promoter in the u11 strain, resulting in a failure to initiate transcription.

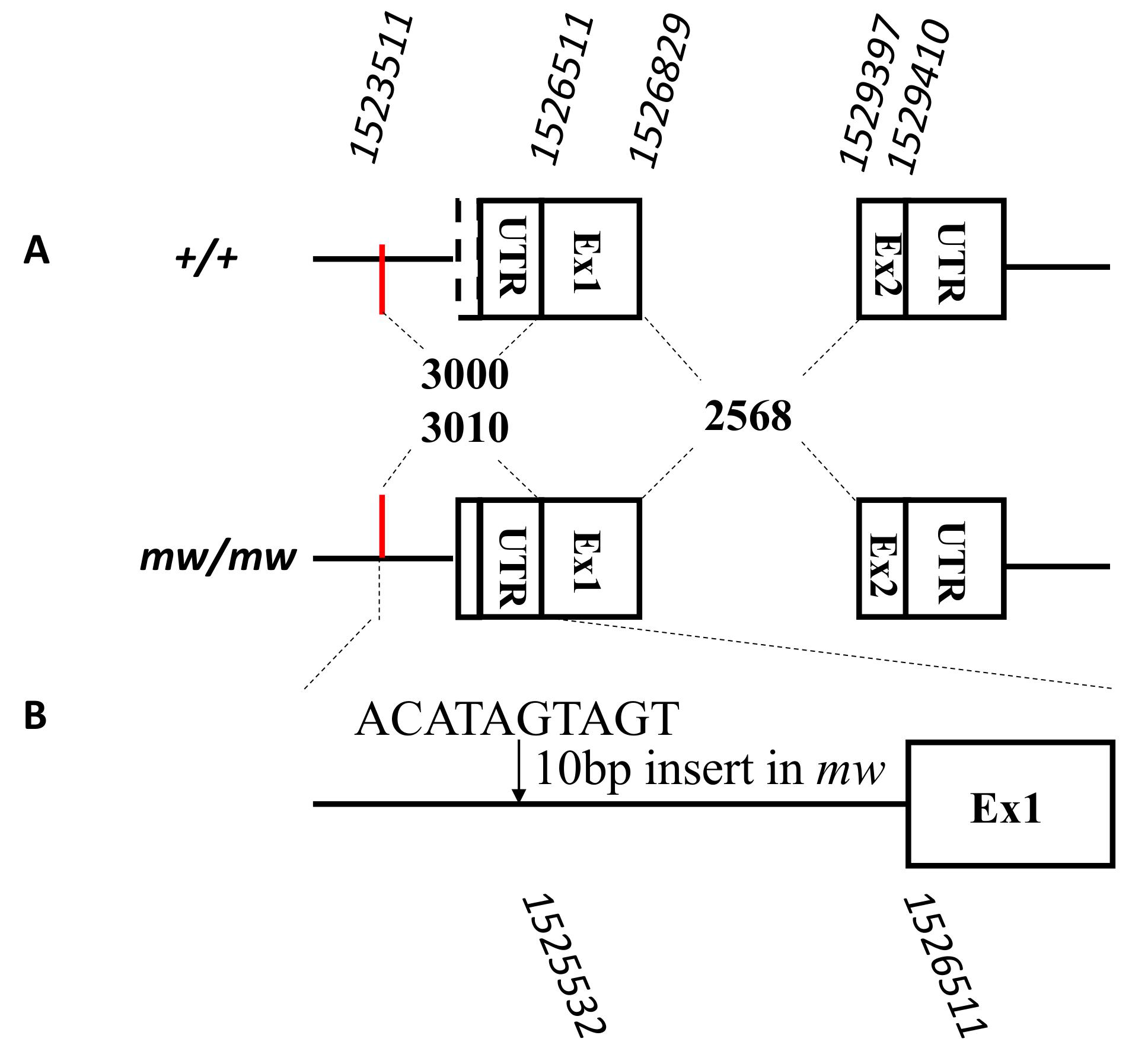
Differences in *m15* between the wild-type, p50, and mutant u11 strains. The two boxes indicate the two exons of *mw* in the p50 and u11 strains. Although they had the same coding sequences, the *mw* mutant strains contained a 10-bp insertion before the initiation codon. The sequence is listed.

### Promoter verification

To verify the promoter activity in the u11 strain, PCR amplification of 1.5 kb of potential *Bm-mw* promoter fragments from the genomic DNA of the u11 and p50 strains was performed (Fig. 6A). As shown in Figure 6A, when compared with the control (p50 wild type), the u11 strain contained an insertion. A cell transfection vector expressing green fluorescent protein named PXLBacII-IE1DsRed2-promoter-EGFP was constructed using a promoter cloned from the different strains (Fig. 6B), and the *Bm-mw* promoter activity was analyzed in the *Bm*-N cell line. The results showed that the *Bm-mw* promoter in the p50 strain had basic transcription activity. However, the promoter from u11 could not initiate the expression of the green fluorescent protein (Fig. 6B), implying that the *Bm-mw* promoter of the u11strain was not induced in response to direct expression of *mw*.

**Figure 6.**
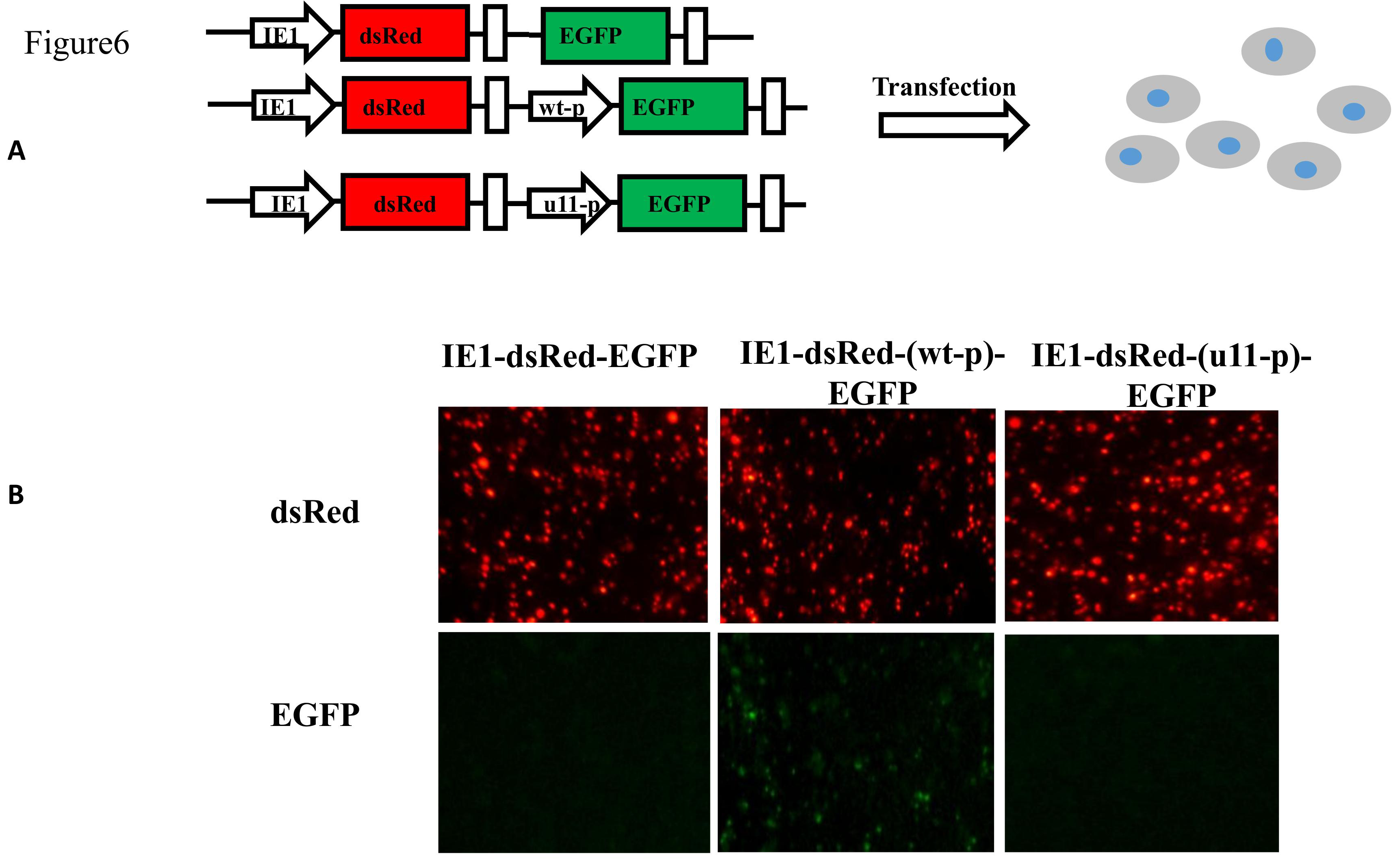
Schematic representation of two transgenic constructs and their validation in *Bm*-N cells. The transgenic pBac-IE1-DsRed-EGFP vector, which contained the full ORF of the *EGFP* reporter gene under the control of the *mw* promoter region in the p50 and u11 strains. A control vector, pBac-EGFP, also contained the *EGFP* ORF region but no promoter region. Both constructs contained the full ORF of the red fluorescent protein DsRed, driven by the IE1 promoter. (B) Validation of two constructs in *Bm*-N cells by fluorescence investigation 96 h after transfection. Red fluorescence was detected in both plasmids of pBac-EGFP, but the reporter gene *EGFP* was expressed only in cells transfected with the pBac-p50 promoter EGFP. DsRed, red fluorescent protein; EGFP, green fluorescent protein.

## Discussion

In insects, the wing is a vital functional organ, which plays an important role in foraging, migration, attracting mates, and evading predators. The shape and size of wings have important functional roles. The adult silkworm, a model lepidopteran insect, retains wings but has lost the ability to fly as a result of artificial breeding, but wing at mating still has a very important role. Not only do studies of genetic mutations enrich understanding of the silkworm mutation gene pool, studies of winged insects can play a positive role in promoting. The *mw* wing mutation is significantly less common than normal-type wings, and information is lacking on its function. Therefore, related genes will help to parse the development of the insect wing control machine system. The information obtained can be applied to the genetic control of lepidopteran pests.

Classical genetic studies revealed mutations in silkworm minute wing Chain group of 22. The evidence in the present study supports the hypothesis that the palmitoyltransferase ZDHHC18-like gene is responsible for the minute wing mutant. First, the gene responsible for the *mw* mutant was genetically mapped within a 275-kb region. Second, the embryonic CRISPR/Cas9 for *Bm-mw* elicited a minute wing phenotype similar to that of the u11 mutant. Third, a *Bm-mw* ortholog in other different species, including *Drosophila melanogaster*, encoded palmitoyltransferase, a member of the DHHC family, which regulates the development of the wing disc. Based on these findings, we conclude that the palmitoyltransferase ZDHHC18-like gene is responsible for the *mw* mutation and essential for the development of wings in the silkworm.

Palmitoyltransferase is responsible for adding palmitates to cytoplasmic proteins (Linder and Deschenes 2007, Nadolski and Linder 2007). APP, the ortholog in *D. melanogaster*(Figure 7B), is a precursor of the amyloid b peptide, which is synthesized as a transmembrane glycoprotein(Rosen, Martin-Morris et al. 1989, Harwell and Coleman 2016). A previous study reported that *Drosophila* amyloid precursor protein binding protein 1 (app-bp1) mutations affected apoptosis, leading to small imaginal discs(Kim, Kim et al. 2007). APP was shown to encode DHHC palmitoyltransferase and regulate Fat signaling and Dachs activity in the Hippo pathway(Matakatsu and Blair 2008). In the present study, *Bm*-mw mRNA was more highly expressed in the integument than in other larval tissues. We performed qRT-PCR analysis of the transcription levels of some pivotal genes of the Hippo pathway in the wing disc of wandering-stage u11 mutants(Figure 8). The results indicated that a number of genes, including Hippo and Yki, in the Hippo pathway were respectively down-regulated and up-regulated in the u11 mutant (Fig. 8). These findings suggested that the *Bm-mw* gene might be located upstream of the Hippo and suppress it.

**Figure 7.**
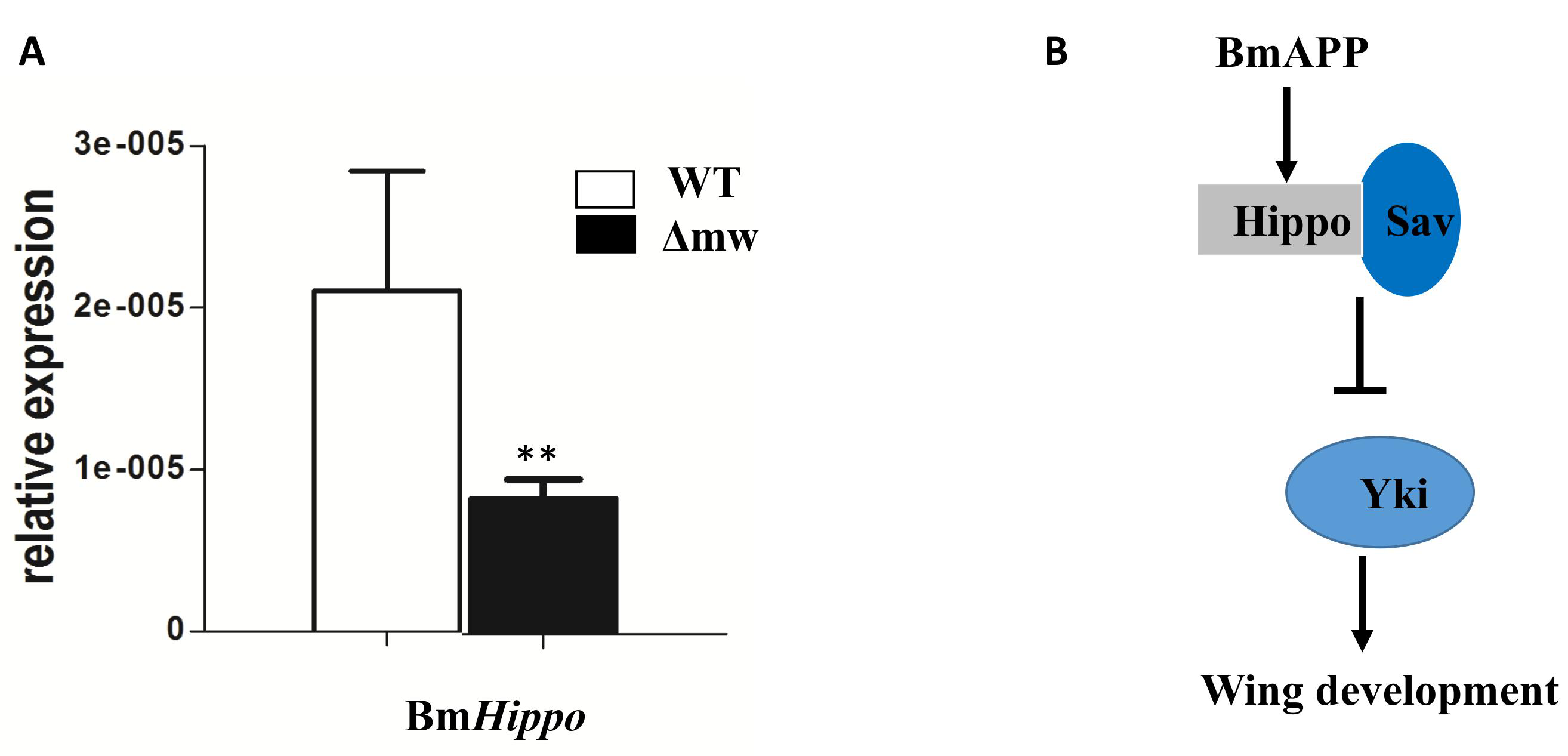
BmHippo is the target of BmAPP. (A)The transcript level of the downstream effector gene, BmHippo, is reduced. The asterisk (**) indicate the significant differences(P < 0.01) compared with the relevant control with a 2-tailed t-test. Error bars depict §SEM. (B) Model showing regulation of BmAPP in B. mori. BmAPP regulated BmHippo which represses BmYki, then affects Hippo signaling leading to the wing development.

In insects, the wing is an important organ, which has underpinned their development, evolution, and survival. Research on genes that affect critical stages in the development of insect wings can shed light on developmental abnormalities in insects, enhance understanding of their classification and evolution, and guide genetic control of insect pests. The silkworm is an ideal model insect for mutation studies because it has been subject to thousands of years of artificial selection. The abundance of silkworm wing-type mutations is an advantage for research of gene function and wing development. Genetic linkage and positional cloning research of the *mw* gene will enrich the silkworm mutation information library. In addition, the regulatory mechanism of insect wing development research has reference the test value. In the future, we will compare wing disc transcriptome differences between p50 and u11 strains, perform gene expression analyses, and analyze genes involved in the pathway. The goal of such studies will be to explore the molecular mechanism underlying the *mw* mutation at a deeper level.

## Materials and methods

### Silkworm strains and cell lines

As *mw* is a recessive single locus, crosses based on a single-pair backcross model were designed (Miao et al., 2005). The u11 strain (*mw/mw)* was crossed with the wild-type p50 strain(+^
*mw*^*/+*^*mw*^). The resulting female F_1_ and male F_1_ crosses were backcrossed with *mw* to generate BC_1_F and BC1M populations, including 22 BC1F individuals (12 were wild type, and 10 were black) and 286 BC1M individuals (144 were wild type, and146 were *mw* mutants). As the silkworm lacks crossing over in females (Maeda, 1939), the BC1F population was used to determine the linkage group and to screen for polymorphisms. A wild-type Nistari strain was used to detect and confirm the genotypes. All the strains, which are preserved in Sericultural Research Institute, Chinese Academy of Agricultural Sciences, were fed fresh mulberry leaves at 25 °C under standard conditions.

The *Bm*-N *B. mori* ovarian cell line was used (Pan, Cai et al. 2010). The cells were maintained in TC-100 medium (AppliChem) containing 10% fetal bovine serum in 25 cm^2^ petri dishes at 27 °C.

### Positional cloning of *mw*

DNA fragment amplification using the polymerase chain reaction (PCR) and the restriction fragment length polymorphism method were used to map the *mw* locus to the genomic DNA sequence. The sequence-tagged site (STS) was searched at various positions along the nucleotide sequence scaffold nscaf3056 on chromosome 22. DNA markers revealed a polymorphism between u11 and p50. The u11 and p50 strains were used for the genetic analysis of BC1 individuals.

### Isolation of genomic DNA and total RNA

Genomic DNA was extracted from adult legs of u11.p50 Nistari and hybrids using a DNA extraction buffer (1:1:2:2.5 ratio of 10% SDS to 5 mol NaCl to 100 mmol EDTA to 500 mmol Tris-HCl, pH = 8), incubated with proteinase K, and then purified via standard phenol: chloroform extraction and isopropanol precipitation extraction, followed by RNaseA treatment. PCR amplification was carried out using 50 ng of genomic DNA as the template. The PCR conditions were as follows: 98 °C for 2 min, 35 cycles of 94 °C for 10 s, 55 °C for 30 s, and 72 °C for 1 min, followed by a final extension period of 72 °C for 10 min. The PCR product was subcloned into a pMD-19T (Takara, Kyoto, Japan) vector and sequenced.

Total RNA was isolated from wing discs using TRIzol reagent (Invitrogen, CA) according to the manufacturer’s instructions and subsequently treated with DNase I (Invitrogen) to remove genomic DNA. A ReverAid First Strand cDNA Synthesis Kit (Fermentas, Lithuania) was used to for cDNA synthesis using 1 g of total RNA.

### RT-PCR

Total RNA was isolated from the wing discs of wandering-stage larvae of strains p50 and u11, as described above. The GeneRacer Kit (Invitrogen) and 5′- and 3′ RACE procedures were used to obtain full-length cDNA. RT-PCR was performed using Ex Taq under the following conditions: initial denaturation at 94 °C for 2 min, 35 cycles of denaturation at 94 °C for 30 s and 57–60 °C for 30 s, and extension at 72 °C fir 10 min. The PCR products were subcloned into a pMD-19T (Takara) vector and sequenced.

### Preparation and microinjection of Cas9/sgRNA

The Cas9-sgRNA system was used to target candidate*mw*. S1 and S2 were identified by screening the candidate *mw* open reading frame (ORF), following the 5’-GG-N18-NGG-3’ rule(Engel and Grimaldi 2004). The sgRNAs were prepared using a MAXIscrip^R^ T7 kit (Ambion), according to the manufacturer’s instructions. Cas9 mRNA was synthesized using an mMESSAGE mMACHINE^R^ T7 kit (Ambion), according to the manufacturer’s instructions, as previously reported(Wang, Li et al. 2013).

Fertilized eggs were prepared as previously described (Wang, Li et al. 2013) and micro-injected within 3 h after oviposition. sgRNA-1 (150 ng/ml) and sgRNA-2 (150 ng/ml) of the candidate gene were combined with Cas9 mRNA (300 ng/ml), mixed, and co-injected into 480 embryos. Injection with an equal amount of PBS was performed as a control. The injected eggs were incubated at 25 °C in a humidified chamber for 10–12 days until larval hatching.

### Plasmid construction and cell transfection

A transgenic plasmid PXLBacII-IE1DsRed2 was constructed to generate pBac-EGFP, as previously described(Xu, Wang et al. 2014). In the transgenic PXLBacII-IE1DsRed2-promoter(u11)-EGFP plasmid, the *EGFP* reporter gene was driven by a putative promoter sequence. A control plasmid, PXLBacII-IE1DsRed2-promoter(p50)-EGFP, was also constructed. The promoter sequence was PCR amplified and introduced into *KpnI* and *ApaI* restriction sites of pBac-EGFP to generate pBac-promoter-EGFP. The pBac-promoter(u11)-EGFP plasmid and pBac-promoter(p50)-EGFP were transfected separately into *Bm*-N cells using lipofectamine reagent (Invitrogen, Carlsbad, CA), following the manufacturer’s protocol. For transfection, 200 ng of each plasmid were transfected into *Bm*-N cells in a 24-well cell culture plate in duplicate. EGFP fluorescence was investigated 72 h after transfection under a fluorescence stereomicroscope (Nikon AZ100, Tokyo, Japan).

## AUTHOR CONTRIBUTIONS

M. Li. conceived the project. Y. Yu. and X. Liu. performed most of the experiments. X. Ma. performed the expriments about STS molecular marker. N. Liu and C. Hou helped prepared for materials. Z. Zhang. helped during the silkworm genetic transformation. Y. Yu. and M. Li. wrote the manuscript.

